# Bovine milk glycoproteins inhibit SARS-CoV-2 and influenza virus co-infection

**DOI:** 10.1101/2023.02.20.529234

**Authors:** Hanjie Yu, Wentian Chen, Jian Shu, Xin Wu, Jia Quan, Hongwei Cheng, Xiaojuan Bao, Di Wu, Xilong Wang, Zheng Li

**Author notes:** Corresponding author: Zheng Li, Laboratory for Functional Glycomics, College of Life Sciences, Northwest University, No. 229 Taibai Beilu, 710069 Xi’an, China. These authors contributed equally: Hanjie Yu and Wentian Chen.

## Abstract

The attachment of S1 subunit of spike (S) protein to angiotensin-converting enzyme 2 (ACE2) is the first and crucial step of SARS-CoV-2 infection. Although S protein and ACE2 are heavily glycosylated, the precise roles of glycans in their interactions are still unclear. Here, we profiled the glycopatterns of S1 subunit of SARS-CoV-2 and ACE2, and found that the galactosylated glycoforms were dominant in both S1 subunit and ACE2. Interestingly, S1 subunit exhibited the property of glycan-binding protein (GBP) and adhered to the ACE2 via binding to the galactosylated glycans on the ACE2. Our earlier findings demonstrated that the sialylated glycoproteins isolated from bovine milk potently inhibit and neutralize viral activity against influenza A virus (IAV). Importantly, we proved further that the galactosylated glycans on isolated glycoproteins bind to the glycan recognition domains of S1 subunit and competitively inhibit binding of S1 subunit to ACE2 and ultimately impede the entry of SARS-CoV-2 pseudovirus into host cells. We provided a potential protein drug that could be multiple simultaneous inhibitor for coronavirus and IAV co-infection.

## Introduction

The COVID-19 pandemic is caused by SARS-CoV-2 and its variants that are genetically closely related to SARS-CoV-1 to threaten human health and public safety (Wang *et al*, 2020; 2. Tao *et al*, 2021; Li *et al*, 2020). Epidemiology data revealed that the SARS-CoV-2 and its variants have resulted in more than 634 million confirmed infected cases and more than 6.5 million deaths worldwide as of Nov 21, 2022 (https://covid19.who.int/). Worryingly, the co-infection of SARS-CoV-2 and influenza A virus (IAV) is associated with increased odds of receiving invasive mechanical ventilation and death (Swets *et al*, 2022). Unfortunately, there is no effective preventive drug for anti-SARS-CoV-2 and IAV co-infection activities yet.

Glycosylation is the most common protein post-translational modification for all kinds of viruses, which not only promotes viral protein folding and subsequent trafficking, but also modulates their interactions with receptors and the following innate and adaptive immune response to affect the host recognition, viral replication, and infectivity (Carbaugh *et al*, 2020; Watanabe *et al*, 2020). Corolla-like spike protein (S protein) is a characteristic of SARS-CoV-2, which is excessively glycosylated to present 22 N-glycosylation sites and 17 O-glycosylation sites (Zhao *et al*, 2020). The glycosylation of S protein extensively affects host recognition, penetration, binding, recycling, and pathogenesis (Tian *et al*, 2021). Despite its variants maintaining an overall similar glycosylation structural conformation, it is also the case that the sites of N-glycosylation on their S protein can happen to shift (Kuo *et al*, 2022). As receptor of SARS-CoV-2, angiotensin-converting enzyme 2 (ACE2) is also a heavily glycosylated protein to have 7 N-glycosylation sites and 2 O-glycosylation sites (Shajahan *et al*, 2021). It is known that the clove homo-trimeric S protein of SARS-CoV-2 binds to ACE2 is crucial for subsequent membrane fusion and virus entry (Jackson *et al*, 2022; Wrapp *et al*, 2020; Yan *et al*, 2020; V’kovski *et al*, 2021). However, the roles of glycans in the infection of SARS-CoV-2 still need further elucidation. Recently studies revealed that the glycans in sterically mask polypeptide epitopes and directly modulate S protein-ACE2 interactions, and the glycosylation of ACE2 contributes substantially to the binding of the virus (Ramírez, *et al*, 2021; Mehdipour *et al*, 2021). The soluble ACE2 could be a therapeutic option to treat SARS-CoV-2 infection (Alhenc-Gelas *et al*, 2020; Krishnamurthy *et al*, 2021).

The intact glycan structure is important for glycoprotein to bind and inhibit pathogens (Bhatia *et al*, 2014). Our earlier findings demonstrated that the glycoproteins with Siaα2-3/6Gal-linked glycans inhibit and neutralize viral activity against IAV (Yu *et al*, 2018; Wang *et al*, 2020). The glycan-binding protein (GBP) also plays a vital role in microbe-host interactions via glycan recognition (Ohtsubo *et al*, 2006). Here we focused our attention on the mechanisms of S protein-ACE2 interactions that bind and/or convert glycan structures for their binding. We demonstrated the precise interactions of glycan structures between S protein and ACE2. Then, we dissected the galactosylated glycoproteins derived from bovine milk, which could block the binding of S protein of SARS-CoV-2 to ACE2 and inhibit the entry of pseudovirus-bearing S protein into host cells.

The discovery of the glycoproteins rich in galactosylated and sialylated glycans can provide valuable strategies to develop a protein drug that could be multiple simultaneous inhibitor for anti-coronavirus and IAV co-infection.

## Results

### Dominant glycan structures of SARS-CoV-2 S1 subunit and ACE2

In the present study, we first investigated glycan structures of S1 subunit of SARS-CoV-2 and ACE2 using antibody-overlay lectin microarrays, respectively (Fig 1A). The acquired images of S1 subunit and ACE2 were shown in Fig 1B and C. There were 26 and 28 lectins to show effective signals (signal noise ratio > 3) in S1 subunit and ACE2, respectively. The averaged normalized fluorescent intensities (NFIs) of each lectin from S1 subunit and ACE2 were summarized as mean values ± SEM in Appendix Table S1. In general, the glycopattern distributions of S1 subunit and ACE2 were indicated as follows: 1) The galactosylated glycans were dominant glycopatterns on both S1 subunit and ACE2 (accounting for 59.82% and 54.10% of the total signals, respectively). 2) The α-1,3 fucosylation and core fucosylation were detected on both of them (accounting for 15.81% and 12.76% of the total signal, respectively). 3) The other types of complex N-glycans were also presented on them both, such as agalactosylated and bisecting GlcNAc N-glycans. 4) The relative abundance of high-mannose types of N-glycans was higher on ACE2 than that on S1 subunit. 5) Only α-2,3 linked sialic acids binder MAL-II showed effective signals on them both (Fig 1D and E).

**Figure 1.**
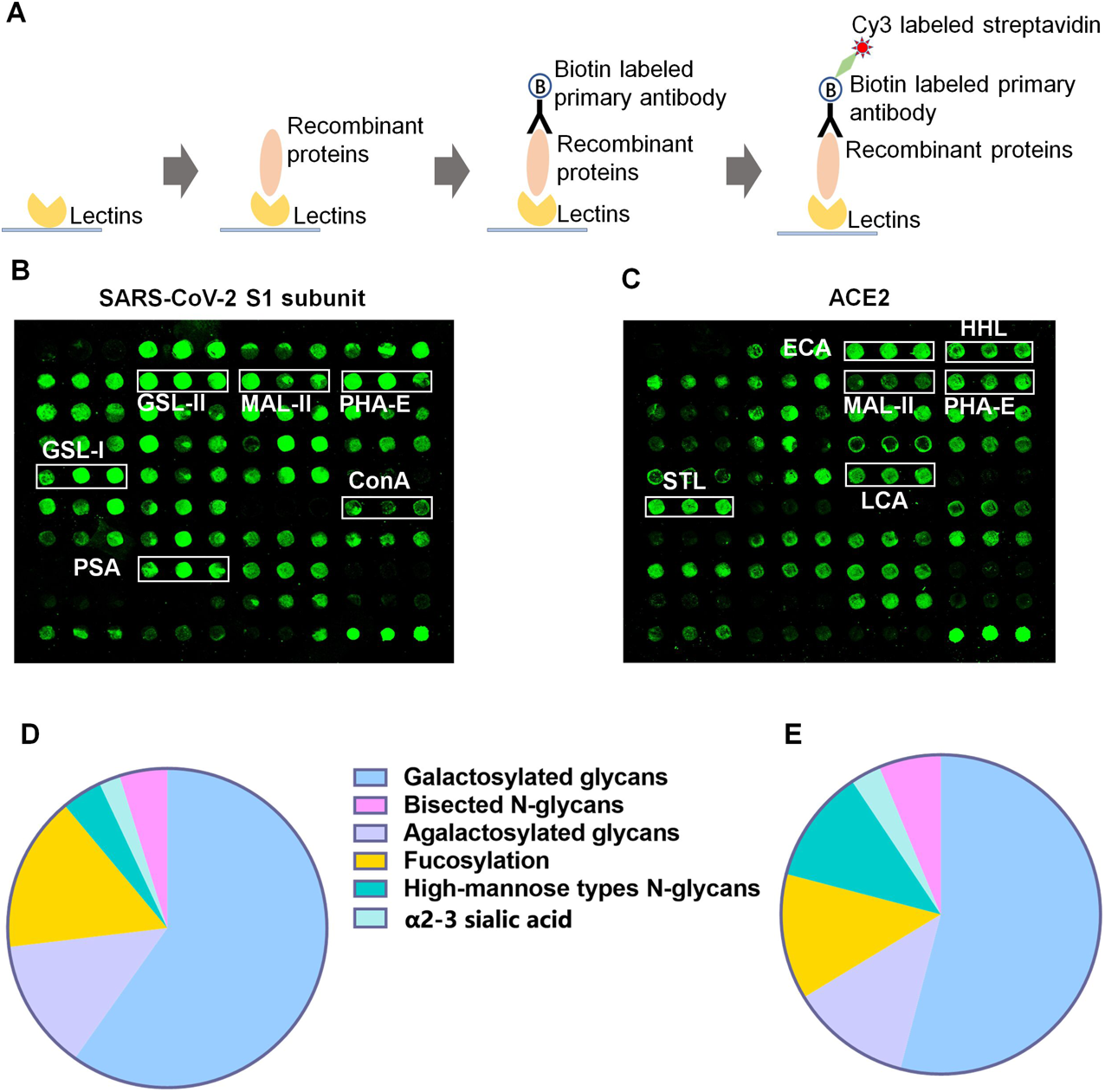
The glycan profiles of SARS-CoV-2-S1 subunit and ACE2 receptor. A The schematic diagram of the preparation of antibody-overlay lectin microarrays. B, C Scanned images were obtained for the analysis of glycopatterns from S1 subunit of SARS-CoV-2 (B) and ACE2 (B). The representative lectins which recognized galactosylated glycans (GSL-I and ECA), agalactosylated glycans (GSL-II and STL), bisected N-glycans (PHA-E), high mannose types of N-glycans (ConA and HHL), fucosylation (PSA and LCA) and α-2,3 linked sialic acid (MAL-II) were marked. D, E The percentage of different types of glycopatterns identified in SARS-CoV-2-S1 subunit (D) and ACE2 (E).

### Twin–twin interaction of SARS-CoV-2 and ACE2

Subsequently, we assessed the roles of N-glycan in the interaction between the S1 subunit of SARS-CoV-2/1 and ACE2 with protein microarrays (Fig 2A). The detection system of protein microarrays was optimized (Appendix Fig S1). Prior to incubation, the PNGase F glycosidase was used to remove N-linked oligosaccharides from S1 subunit of SARS-CoV-2 and SARS-CoV-1 as well as ACE2, then, the interaction between S1 subunit and ACE2 was assessed by protein microarrays. As a result, the binding signals were the highest as ACE2 binds to S1 subunit of SARS-CoV-2 and SARS-CoV-1 in their original glycosylated states, respectively. However, the ACE2 significantly decreased its binding to S1 subunit of SARS-CoV-2 and SARS-CoV-1 in at least one of their de-N-glycosylated states (Fig 2B and C). Interestingly, the binding signals were reduced by 76.08% as ACE2 and S1 subunit of SARS-CoV-2 in their de-N-glycosylated states. The binding signals were almost equivalent as the de-N-glycosylated ACE2 binds to the intact S1 subunit (reduce by 67.62%) of SARS-CoV-2 and the intact ACE2 binds to the de-N-glycosylated S1 subunit of SARS-CoV-2 (reduce by 65.93%). These results indicated that the N-glycans are vital for the binding of S1 subunit and ACE2.

**Figure 2.**
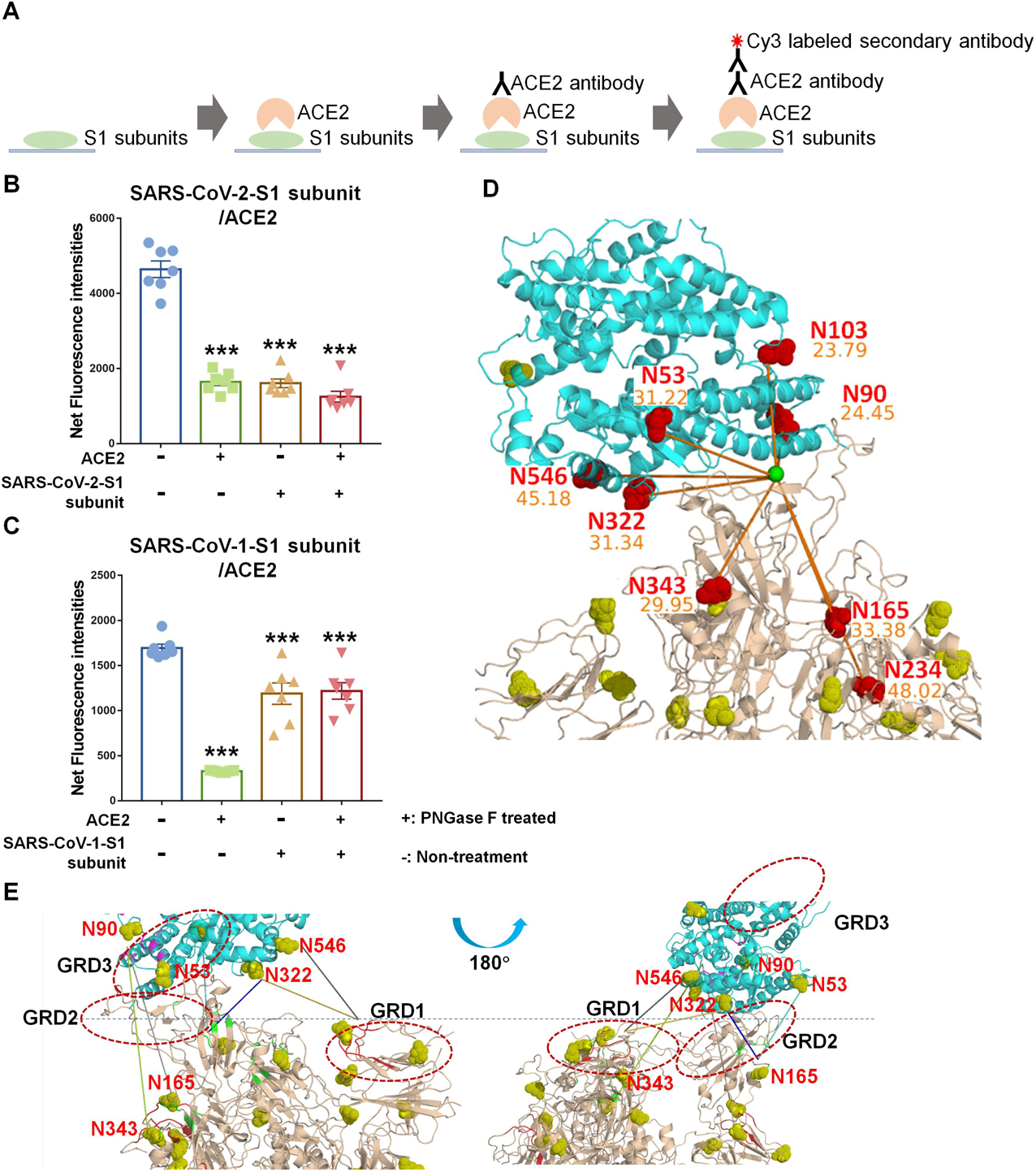
Evaluation for the role of N-glycans in the interaction between S1 subunit and ACE2. A The schematic diagram of the manufacture of SRAS-CoV-2 related recombinant protein microarrays. B, C The N-glycans of S1 subunits of SRAS-CoV-2/1 and ACE2 were removed by PNGase F glycosidase respectively, and the roles of N-glycans in the interaction between SRAS-CoV-1-S1 subunit/ACE2 (B) and SRAS-CoV-2-S1 subunit/ACE2 (C) were evaluated by protein microarrays. The net fluorescence intensities were obtained by subtraction of background and then compared with control by one-way analysis of variance (ANOVA). Data were represented as means ± standard error of mean (SEM). *: p < 0.05, **: p < 0.01, and ***: p < 0.001. D The MD simulation of ithe nteraction of trimeric S protein and ACE2. The N-glycosites on ACE2 and S1 subunit with the distance to the center of the binding interface (green globule) less than 50 Å were marked with red spheres, other N-glycosites were marked with yellow spheres. E The interaction of glycans on certain sites and GRDs (marked with red frame) may involve in the binding of S-protein to ACE2.

Since the binding of S1 subunit of SARS-CoV-2 and ACE2 is crucial for the infection of coronavirus. The S1 subunit forms the globular head containing N-terminal domain (NTD), receptor binding domain (RBD) and smaller subdomains. We investigated further the roles of N-glycosylation sites in the interaction between trimeric S protein and ACE2 with molecular dynamics simulation. There are five sites (N53, N90, N103, N322 and N546) of ACE2 and three sites of S1 subunit (N343 of RBD, N165 and N234 of NTD) distribute surround the interactive interface (<50 Å to the binding center), and most of these N-glycosylation sites (excluding N546 of ACE2) with the distance shorter than 30 Å, which implied the N-glycans on these sites are likely involved in the interaction of S protein and ACE2 (Fig 2D and E). We found further that there are three potential glycan recognition domains (GRDs) in SARS-CoV-2-S1 subunit (GRD1 and GRD2), as well as ACE2 (GRD3), which may interact with the glycans on the above-mentioned sites of S1 subunit and ACE2. Further, molecular dynamics (MD) simulation was used to predict the average distance among glycans on these sites and GRDs. As a result, the glycans on N546 and N332 sites of ACE2 may interact with GRD1 of S1 subunit, the glycans on N53 and N322 sites of ACE2 are very likely to interplay with GRD2 of S1 subunit (average distance < 15Å). In addition, the glycans on N165 and N343 sites of S1 subunit are probable to interact with GRD3 of ACE2 (average distance is approximately 30Å) (Fig EV1).

The above results indicated the glycans are pivotal for S1 subunit of SARS-CoV-2 bound with ACE2, and there is a twin-twin interaction between them, which could be GBPs and interact with glycans on each other. Our results revealed that S1 subunit exhibits higher affinities to glycans than ACE2, meanwhile, the distances between glycans on ACE2 to GRDs of S1 subunit are less than < 15Å, which is shorter than the distances between glycans on S1 subunit to GRD of ACE2. These findings suggest S1 subunit prefers to be a GBP and interacts with glycans on ACE2.

### Carbohydrates for blocking of SARS-CoV-2 and ACE2 binding

The molecular docking analysis was used to predict the potential binding capacity of S1 subunit of SARS-CoV-2 and its variants as well as ACE2 to different carbohydrates. Compared to ACE2, the NTD and S1 subunit of SARS-CoV-2 and its variants (Delta and Omicron) exhibit high affinities to galactosylated structures (Galβ1-3/4GlcNAc, Galβ1-3GalNAc, and GlcNAcβ1-6(Galβ1-3)GalNAc). Interestingly, S1 subunits of both SARS-CoV-2 and its variants show high affinities to galactosylated structures, which implies that mutations of SARS-CoV-2 do not impact the affinities of S1 subunit to the galactosylated structures (Fig 3A).

**Figure 3.**
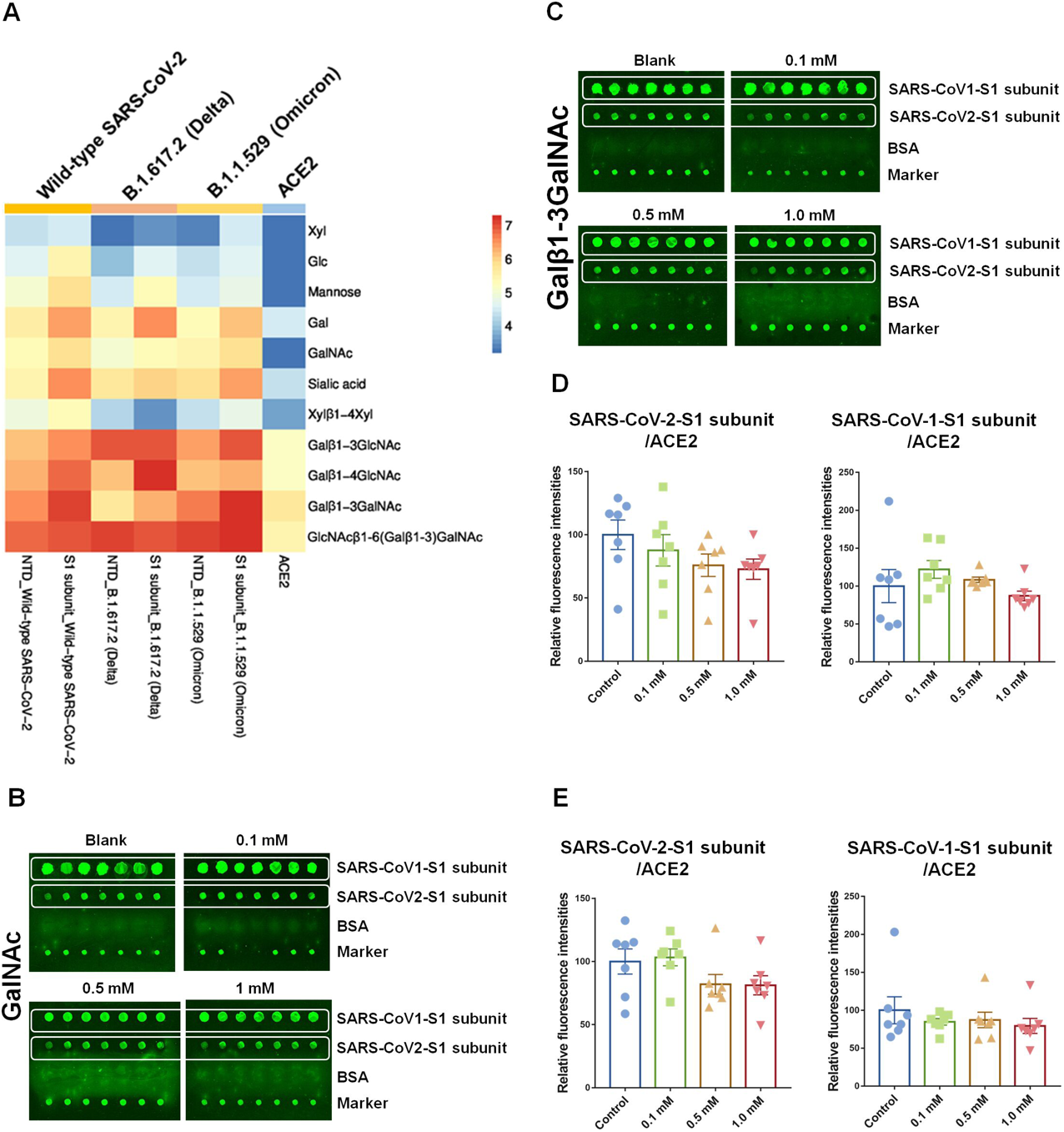
Evaluation of Carbohydrates for blocking of SARS-CoV-2 and ACE2 binding. A A series of carbohydrates for their binding capacity with S1 subunit of SARS-CoV-2 (wild type) and its variants (Delta and Omicron) as well as ACE2 by molecular docking analysis, respectively. The carbohydrates were listed in columns, the NTD, S1 subunit, and ACE2 were listed in rows, and the different affinities were represented by the absolute values of the binding free energy, which were marked by the color of each square, red: high affinity, mazarine: low affinity. B, C Scanned images of protein microarrays. ACE2 (1μg) was mixed with 0, 0.1, 0.5, and 1.0 mM of GalNAc (B) or Galβ-1,3GalNAc (C) and applied to protein microarrays, respectively. D, E Effect of GalNAc (D) and Galβ-1,3GalNAc (e) on the interaction of S1 subunit of SARS-CoV-2/1 and ACE2. The binding signals were extracted and the relative fluorescence intensities were compared with controls by One-Way ANOVA. Data were represented as means ± SEM (error bars). *: p < 0.05, **: p < 0.01, and ***: p < 0.001.

To explore whether free saccharides impact the interaction of S1 subunit of SARS-CoV-2/1 with its receptor ACE2, the different concentrations (0, 0.1, 0.5, and 1.0 mM) of the GalNAc and Galβ1-4GalNAc were added into incubation solution and applied to protein microarrays, respectively (Fig 3B and C). Interestingly, the interaction between S1 subunit of SARS-CoV-2/1 and ACE2 did not evidently interrupt by GalNAc (Fig 3D). However, the binding signals of S1 subunit of SARS-CoV-2 and ACE2 were slightly interrupted by 1.0 mM of Galβ1-3GalNAc (reduced by 18.86%, p >0.05) (Fig 3E).

### Glycoproteins for inhibition of SARS-CoV-2 and ACE2 binding

We have analyzed the glycan structures of the isolated glycoproteins from bovine milk by a lectin microarray (Wang *et al*, 2020) (Fig 4A). The glycopatterns of the isolated glycoproteins were similar to the ACE2, such the galactosylated glycans are dominant (accounting for 52.19% of the total signal) but except for Siaα2-6Gal-linked structures (Fig 4B). Then, we evaluated whether the isolated glycoproteins could serve as competitive multivalent substrates to interfere binding of S1 subunit of SARS-CoV-2/1 with ACE2 by using protein microarrays. We observed that the binding signals of S1 subunit of SARS-CoV-2/ACE2 are significantly reduced as 0.21-0.83 ng/μL of the isolated glycoproteins were added (*p* <0.001). Meanwhile, the interaction of SARS-CoV-1-S1 subunit and ACE2 was moderately disturbed by 0.04 and 0.08 ng/μL of the isolated glycoproteins, and was remarkably interrupted as 0.21-0.83 ng/μL of the isolated glycoproteins were added (*p* <0.001) (Fig 4C). These results revealed that the isolated glycoproteins with abundant Gal and Siaα2-3/6Gal-linked glycan structures from bovine milk could serve as competitive substrates and interfere with the interaction of S1 subunit of SARS-CoV-2/1 with its receptor ACE2.

**Figure 4.**
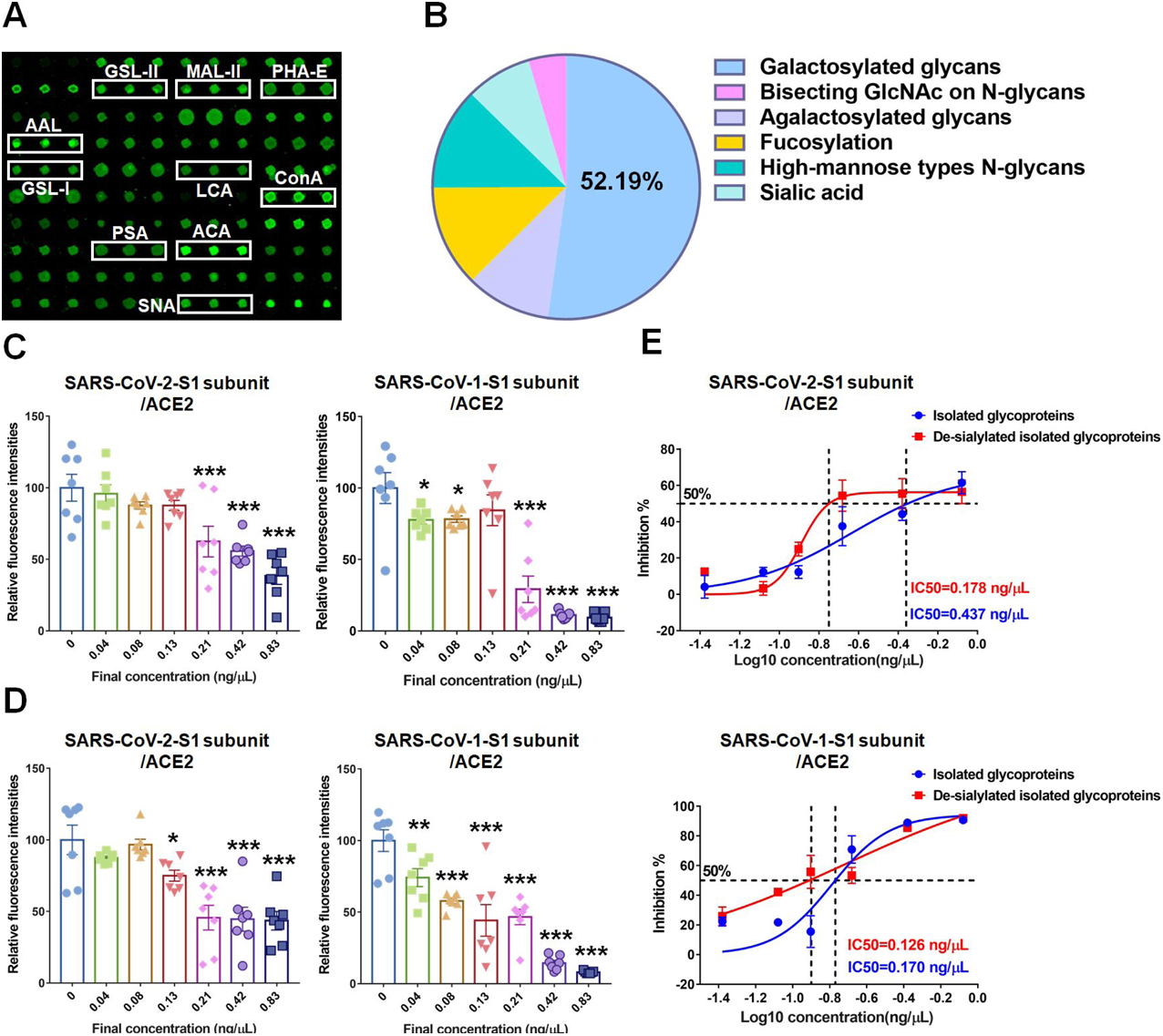
The isolated glycoproteins for inhibition of SARS-CoV-2 and ACE2 binding. A The scanned image was obtained from the lectin microarray analysis of glycoproteins isolated from bovine milk. The representative lectins which recognize galactosylated glycans (ACA and GSL-I), agalactosylated glycans (GSL-II), bisected N-glycans (PHA-E), high mannose types of N-glycans (ConA), fucosylation (AAL, PSA and LCA), α-2,3 linked sialic acid (MAL-II) and α-2,6 linked sialic acid (SNA) were marked with white frames. B The percentage of different types of glycopatterns identified in the isolated glycoproteins. C, D The evaluation of the effect of intact and de-sialylated isolated glycoproteins on interaction of S1 subunit and ACE2. 0, 0.04, 0.08, 0.13, 0.21, 0.42 and 0.83 ng/μL of intact isolated glycoproteins (C) or de-sialylated isolated glycoproteins (D) were mixed with 1 μg of ACE2 and incubated with protein microarrays, respectively. The relative binding intensities of each group were compared with control, the significant difference between groups was determined by One-Way ANOVA. Data were represented as means ± SEM (error bars). E Inhibition curves for intact isolated glycoproteins (upper) and de-sialylated isolated glycoproteins (lower). The four-parameter inhibition curves are generated and the particular IC50 value of intact isolated glycoproteins and de-sialylated isolated glycoproteins in this graph are indicated. Data are represented as means ± SEM (error bars), *: p < 0.05, **: p < 0.01, and ***: p < 0.001.

To determine whether sialic acids of isolated glycoproteins impact the interaction of S1 subunit and ACE2, the sialic acid moieties of the isolated glycoproteins were released by α2-3/6/8 neuraminidase. The abundance of galactosylation was evidently increased in isolated glycoproteins treated with neuraminidase (Appendix Fig S2). Compared to intact glycoproteins, the binding of S1 subunit of SARS-CoV-2 and ACE2 were moderately disturbed by 0.13 ng/μL of the de-sialylated glycoproteins (*p* <0.05). Similarly, the binding of S1 subunit of SARS-CoV-1 and ACE2 was remarkably inhibited by 0.04-0.83 ng/μL of the de-sialylated glycoproteins (*p*<0.01) (Fig 4D). Moreover, the de-sialylated isolated glycoproteins potently inhibited the bindings of S1 subunit of SARS-CoV-2/1 and ACE2 with 50% inhibitory concentration (IC50) of 0.178 ng/μL and 0.126 ng/μL, respectively, which were lower than that of intact isolated glycoproteins. However, the inhibition effects did not show an obvious difference between intact and de-sialylated glycoproteins at high concentrations (more than 0.208 ng/μL) (Fig 4E). These data indicated that galactosylated glycans but not sialylated glycans of isolated glycoproteins disturb the interaction of S1 subunit and ACE2.

### Glycoproteins for inhibition of SARS-CoV-2 pseudovirus into host cells

Subsequently, we utilized a replication-competent pseudovirus that carries the SARS-CoV-2 S protein as a surrogate model to evaluate the inhibitory effect of the isolated glycoproteins for entry of SARS-CoV-2 into host cells (Li *et al*, 2020). The specific protein bands of S protein (S1 + S2) and S1 subunit were observed at positions of about 200 kDa and 110 kDa in cell lysis and concentrated culture supernatant. In addition, the dramatic cytopathic effects (CPE) and visible plaques were observed in Vero E6 cells after 72 h post-infection (Fig EV2A-C). These results indicated the S protein-bearing pseudovirus effectively infects host cells. Pseudovirus-based CPE inhibition assays demonstrated that the isolated glycoproteins prevent pseudovirus-induced CPE in a dose-dependent manner and the 50% inhibitory concentration (IC50) of the isolated glycoproteins is 0.13 μg/μL, however, the equal amount of raw milk proteins and BSA could not inhibit CPE obviously (Fig 5A).

**Figure 5.**
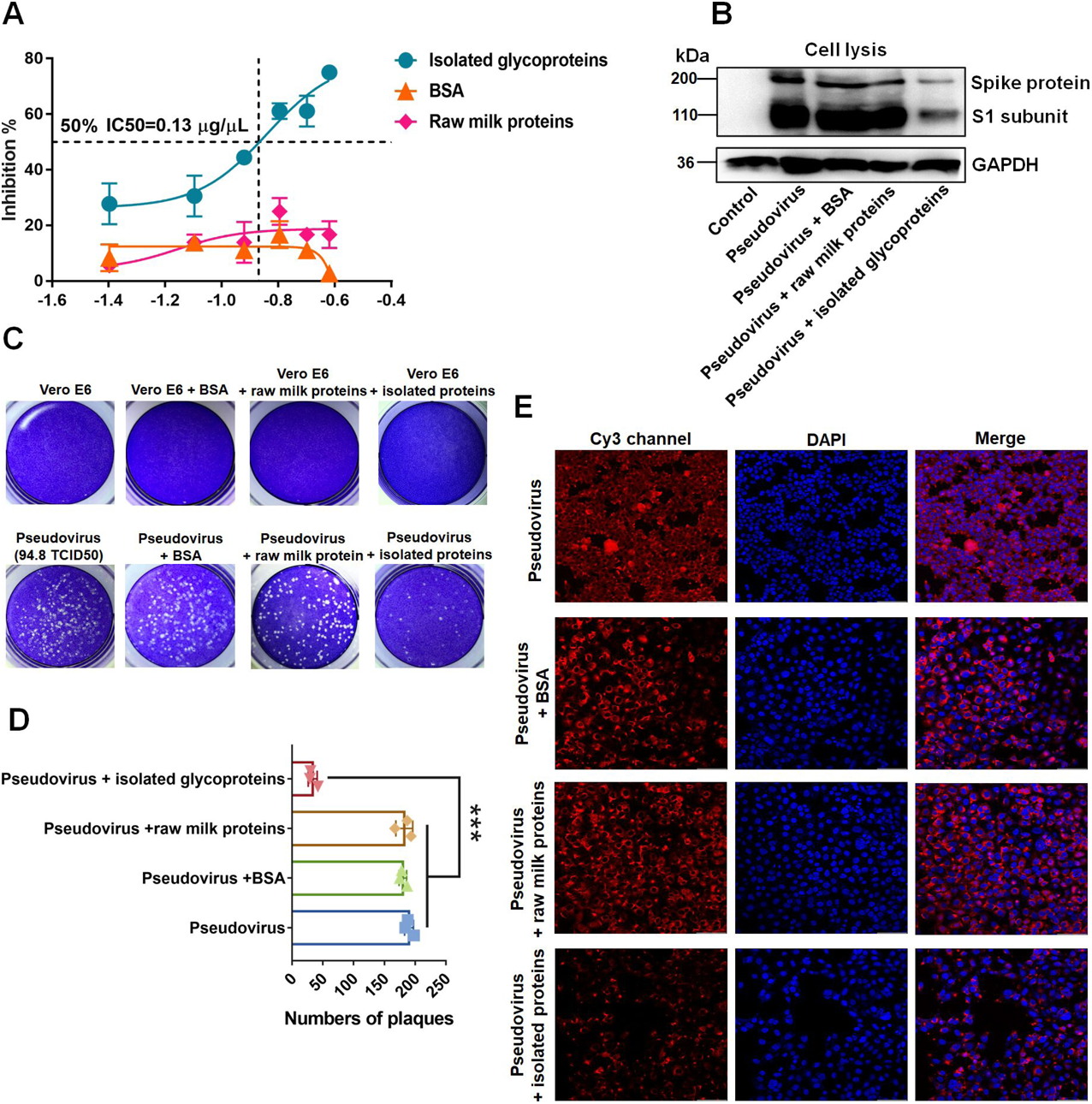
The isolated glycoproteins for inhibition of entry and infection of SARS-CoV-2 pseudovirus to host cells. A Assessment the inhibitory effect of the isolated glycoproteins by pseudovirus-based CPE inhibition assays. Vero E6 cells were seeded into 96 well plates (1×10^4^ cells per well). The pseudovirus (31.6 TCID50 per well) was per-treated with isolated glycoproteins, BSA, and raw milk proteins for 1 h at a concentration ranging from 0.04 to 0.24 μg/μL, and added into plates. The CPE was determined by microscope, and the percentages of inhibition were counted from triplicate plates. The four-parameter inhibition curves were generated and the particular IC50 value of isolated glycoproteins in this graph is indicated. Data were represented as means ± SEM. B The isolated glycoproteins significantly inhibited entry of pseudovirus. Vero E6 cells were inoculated into six-well plates (4×10^5^ cells per well). Pseudovirus (94.8 TCID50 per well) pre-incubated with isolated glycoproteins, BSA and raw milk proteins (final concentration is 0.13 μg/μL) and added into plates. The total proteins were extracted and the expression level of S protein was analyzed by West blotting 48 h after infection. C The isolated glycoproteins significantly suppressed the formation of plaque. Vero E6 cells were inoculated into six-well plates (4×10^5^ cells per well) and incubated as described above. The cells were cultured in 2% FBS/ 1% avicel DMEM medium for 72 h until the plaque was observed. D The statistic results of the number of plaques from triplicate plaque assays. The cells were immobilized and stained, the number of plaques in each well was counted and compared based on groups by using One-Way ANOVA. Data were represented as means ± SEM. *: p < 0.05, **: p < 0.01, and ***: p < 0.001. E Immunofluorescence analysis of the attachment of pseudovirus by confocal microscopy. hACE2/293A cells (3 × 10^4^ cells) were grown on 30 mm confocal dishes and infected with pseudovirus that pre-incubates with isolated glycoproteins, BSA and raw milk proteins as described above. After 6 h incubation, the cells were immobilized, blocked and incubated with primary antibody against SARS-CoV-2 S protein and Cy3 labeled second antibody. Nuclei were counterstained with DAPI. The images were acquired at the same exposure time and shown on the same scale. Bar = 100 μm.

We further investigated the inhibitory effect of the isolated glycoproteins for the entry of SARS-CoV-2 pseudovirus into host cells. SARS-CoV-2 pseudovirus was pre-incubated with 0.13 μg/μL of the isolated glycoproteins, raw milk proteins as well as BSA at 37°C for 1 h, respectively. Then SARS-CoV-2 pseudovirus-protein mixtures were added to Vero E6 cells to test their effect on pseudovirus infection. Our results showed that the expression level of S protein and formation of plaques were significantly suppressed in Vero E6 cells infected by the isolated glycoproteins pre-treated pseudovirus pre-treated compared to others (Fig 5B-D). We also assessed the effect of the isolated glycoproteins on the binding of SARS-CoV-2 pseudovirus to 293A-hACE2 cells (Fig EV2D). The results showed a weakened signal of S protein of the pseudovirus in 293A-hACE2 cells infected by the pseudovirus pre-treated with 0.13 μg/μL of the isolated glycoproteins (Fig 5E). Our results indicated that the galactosylated glycoproteins effectively block the entry of SARS-CoV-2 pseudovirus into host cells.

## Discussion

It is well known that the main strategy for the design of inhibitor is blocking the binding of coronavirus to ACE2 receptor on host cells. Several protein and polypeptide inhibitors such as soluble ACE2 and peptide analogues of ACE2 and RBD have been designed based on this principle (Xiu *et al*, 2020; Planas *et al*, 2021). However, the mutations in SARS-CoV-2 are resulting in both amino acid sequences and glycosylation changes, which distinctly impairs the sensitivity and neutralizing activity of antibodies (Sievers *et al*, 2022; Kuo *et al*, 2022; Reis *et al*, 2021). Hence, a new strategy for the design of inhibitor is to disturb the interaction of S protein and ACE2 mediated by glycans. It is known that most of N-glycans on S1 subunit are complex type of N-glycans with Gal terminal (Zhao *et al*, 2020). Several studies reported that erectogenic GBP such as Galectin-3, Galectin-9, and plant lectins strongly bind to glycans on SARS-CoV-2 and disturb the binding of SARS-CoV-2 (Caniglia *et al*, 2020; Aminpour *et al*, 2021; Hoffmann *et al*, 2021; Wang *et al*, 2021). Here, we demonstrated that the complex types of N-glycan are dominant in S1 subunit and ACE2. Moreover, both of them contain a high level of galactosylation, which takes up over 50% of total glycan abundance. As glycoproteins, S protein and ACE2 also exhibited potential binding capacity to glycans, especially S1 subunit of S protein displayed high affinities to galactosylated glycans. Moreover, the glycans on N53, N322, and N546 of ACE2 were highly possible to interact with GRDs of S1 subunit. These findings suggested that S protein prefers to serve as a GBP and adhere to the galactosylated glycans on ACE2.

Emerging SARS-CoV-2 variants, such as Delta and Omicron have impacts on viral transmissibility and pathogenicity as well as vaccine effectiveness (Kumar *et al*, 2022). Delta variants have grown dominant internationally and are still evolving. Omicron variants contain numerous mutations that influence their behavior (Haveri *et al*, 2022; Benning *et al*, 2022). We found that the S1 subunits of original strain and variants indicate higher affinities to galactosylated glycans than other carbohydrates. These findings implied that galactosylated glycans conjugated with proteins could be a potential inhibitor against Delta and Omicron variants.

Recent studies have shown that the galectin-like NTD of S1 subunit mediates SARS-CoV-2 interaction with sialic acids, contributing to SARS-CoV-2 adhesion (Reis *et al*, 2021; Caniglia *et al*, 2020). The interaction between the NTD of S1 subunit and sialic acids on the surface of host cells may be critical for SARS-CoV-2 cell entry (Sun, 2021). However, the studies by Yang *et al* reported that sialidase treated of S protein and ACE2 does not markedly impact viral entry (Yang *et al*, 2020). Our findings indicated that the inhibitory effect of de-sialylated isolated glycoproteins (higher level of galactosylation) on the binding of S1 subunit to ACE2 is greater than intact isolated glycoproteins (contained sialylation and galactosylation). It indicated that the galactosylated but not sialylated glycans on isolated glycoproteins competitively inhibit the binding of S1 subunit to ACE2.

There were mounting concerns that respiratory virus co-infections are more likely to occur during future winters, and the clinical outcome of respiratory viral co-infections with SARS-CoV-2 is unknown (Swets *et al*, 2022). The studies by Bai *et al* reported that IAV has a unique ability to aggravate SARS-CoV-2 infection, increase SARS-CoV-2 viral load and more severe lung damage in mice coinfected with IAV, therefore, prevention of IAV infection is of great significance during the COVID-19 pandemic (Bai *et al*, 2021). Here, our earlier studies have demonstrated that the isolated glycoproteins containing a high abundance of Siaα2-3/6Gal-linked glycans, could inhibit and neutralize viral activity against IAV (Yu *et al*, 2018; Wang *et al*, 2020). We proved further that the galactosylated glycans on the isolated glycoproteins could also significantly inhibit the binding of S1 subunit of SARS-CoV-2/1 to ACE2 and impede the binding and entry of SARS-CoV-2 pseudovirus.

We concluded that the protein portion of the isolated glycoproteins serves as a steady multifunctional scaffold, the special galactosylated and sialylated glycans on the proteins interact with the S1 subunit of coronaviruses and HA of influenza viruses and competitively inhibit the bindings of viruses to glycans on receptors and host cells. Our study takes key insights into the first step of coronaviral attachment and provided a potential protein drug that could be a multiple simultaneous inhibitor for coronavirus and influenza virus co-infection.

## Materials and Methods

### Analysis of antibody-overlay Lectin Microarrays

The lectin microarrays were produced as described previously (Yu *et al*, 2019; Liu *et al*, 2018). Briefly, 37 lectins with different binding preferences covering N- and O-linked glycan structures were dissolved in recommended buffers according to the manufacturer’s instructions and spotted on homemade epoxysilane-coated slides using a smart arrayer 48 (Capitalbio, Beijing, China). Each lectin was spotted in triplicate per block, with quadruplicate blocks on one slide.

The antibody-overlay lectin microarray assays were performed as described previously with minor modifications (Meany *et al*, 2011). Firstly, lectin microarrays were blocked with blocking solution (2% (*w/v*) BSA in PBST) for 1 h, then 1 μg of recombinant proteins of SARS-CoV-2-spike protein S1 subunit and ACE2 (expressed by HEK293 cell, purchase form acrobiosystems, China) were diluted with PBS-T (containing 1% (*v/v*) Triton X-100) and applied to lectin microarrays and incubated overnight at 25℃, respectively. After that, 20 μg of mouse IgG (Bioss, China) was used to re-block for 1 h to eliminate the binding of lectins with the primary antibodies. Then, 200 ng of anti-SARS-CoV-2-spike protein S1 subunit or anti-ACE2 biotinylated primary antibodies (Bioss, China) was applied to microarrays. After that, 200 ng of Cy3 labeled streptavidin (homemade) was incubated with lectin microarrays. The microarrays were washed and scanned using a Genepix 4000B confocal scanner (Axon Instruments, Foster City, Calif., USA). The acquired images were analyzed at 532 nm for Cy3 detection by Genepix 7.0 software.

### Molecular docking analysis of the interaction between SARS-CoV-2 variants and different carbohydrates

Due to the partial peripheral elements lack of the available S-protein structure, the whole amino acid sequence from one S protein (Genebank ID: QHN73810.1) was selected for homologous modeling by SWISS-MODEL web service (Chen *et al*, 2012). The 3D structure file of S protein with the largest similarity of 99.9 % to a reported S protein of SARS-CoV-2 (PDB ID: 6VSB) was created. For structural optimization, 100 ns of MD simulation was adopted. Minimization and equilibration were performed using the NAMD 2.8 program with the CHARMM22 all-atom force field for the protein as described previously (Trott *et al*, 2010).

Subsequently, a series of carbohydrates (including monosaccharides and oligosaccharides) were designed to assess the potential glycan-binding ability of S1 subunit and NTD of wild type of SARS-CoV-2 and its variants (Delta (PDB ID: 7W92), Omicron (PDB ID:7WVO)) and ACE2 receptor (PDB ID: 5W9J). The different carbohydrates were constructed by the online SWEET-Ⅱ program and optimized by the MM3 force field (Lemmin *et al*, 2019). The automated docking analysis between proteins and carbohydrates was performed by using the AutoDock Vina program. A grid box with the size of 120 × 120 × 120 points grid box was used to cover these proteins during the docking analysis. The receptor atom positions were held fixed, and the glycosidic bonds of glycans were flexible. Other docking parameters were set to default (Turoňová *et al*, 2020).

### Molecular Dynamic Simulation for glycosylated S-protein and ACE2 complex

To investigate the roles of N-glycans in the interaction of S protein and ACE2, the MD simulation of glycosylated S protein-ACE2 complex was performed based on the reported structure file (PDB ID: 6VSB). The “complex” type N-glycans were added at all N-glycosylated sites by “glycoprotein builder” program in GLYCAM online server. The NVT ensemble and a revised force-field file for glycoprotein were used in the MD simulation for these complexes, and further distance analysis for the N-glycans was adopted by VMD1.9.3. Six pairs of distances between the geometric centers of three GRDs and the terminal monosaccharide on the nearby N-glycosylation sites were selected for potential interaction assessment. The distance of “GRD1-N90, GRD1-GlcNAc, GRD1-MAN, GRD1-Gal, and GRD1-SA” were sampled every 100 ps during 100 ns MD simulation.

### Protein microarrays

The manufacture of protein microarray was similar to lectin microarray. In brief, the recombinant proteins (including S1 subunit of SARS-CoV-2 and SARS-CoV-1) were dissolved into PBST (PBS containing 0.01% (*v/v*) Tween-20) at the same concentration, and spotted onto epoxysilane-coated slides. BSA and Cy3 labeled BSA served as negative control and position marker, respectively. Each sample was spotted seven times per block. The protein microarray was blocked and then incubated with 1 μg of ACE2 overnight at 25 ℃. Subsequently, 200 ng of anti-ACE2 primary antibody (mouse monoclonal, proteintech, China) and Cy3 labeled goat against mouse second antibody (Bioss, China) were applied to microarrays. The acquired images were obtained by Genepix 4000B confocal scanner and analyzed at 532 nm for Cy3 detection by Genepix 7.0 software.

### Treatment of glycosidases

The N-glycans of ACE2 were removed by PNGase F glycosidase (NEB) according to the manufacturer’s instructions. To remove N-glycans of S1 subunits of SARS-CoV2/1 on protein microarray, 1 μL of PNGase F glycosidase was added into 120 μL 1× GlycoBuffer (with 0.01% (v/v) NP-40) and incubated with protein microarrays at 37°C for 1 h to hydrolyze the N-glycans on S1 subunits. Then, the protein microarrays were washed and dried. The effect of N-glycans on the interaction of S1 subunit and ACE2 was assessed by protein microarrays as described above.

### Assessment of the effect of GalNAc and Galβ1-4GalNAc on the interaction of S1 subunit and ACE2

The inhibition effect of GalNAc and Galβ1-4GalNAc were assessed by the protein microarrays. Briefly, 0, 0.1, 0.5 and 1.0 mM Gal (Sigma) or Galβ1-4GalNAc (Bio-get, China) was added into incubation solution (contained 1 μg of ACE2), respectively, and incubated with the protein microarrays overnight at 25℃. After washed with PBST and PBS, protein microarrays were incubated with antibody against ACE2 and Cy3 labeled second antibody, respectively. The protein microarrays were scanned and fluorescence intensities were extracted by Genepix 7.0 software.

### Assessment of the inhibition effect of isolated glycoproteins on the interaction of S1 subunit and ACE2

The glycoproteins were isolated from bovine milk as described (Wang *et al*, 2020). To obtain de-sialylated glycoproteins, the isolated glycoproteins are treated by α-2,3/6/8 neuraminidase (NEB) according to the manufacturer’s instructions. The levels of sialylation and galactosylation in intact and de-sialylated isolated glycoproteins were determined by lectin blotting. The inhibitory effect of intact and de-sialylated isolated glycoproteins on the interaction of SARS-CoV-2/1-S1 subunits and ACE2 were also assessed by the protein microarrays.

### Cell Culture and Cell Line Construction

Human embryonic kidney 293A (293A) and Vero E6 cells were maintained in Dulbecco’s modified Eagle’s medium (DMEM, Corning, USA) supplemented with 10% (*v/v*) fetal bovine serum (FBS, Newzurm, New Zealand) and 1% (*v/v*) Antibiotic-Antifungal (MCE, China). To overexpress ACE2, 293A cells were transfected with the plasmid of pCDNA3.1-ACE2 (human)-3×FLAG (MiaoLing Plasmid Platform, China) using polyfect (QIAGEN, West Sussex, UK) according to the manufacturer’s protocol. Cells were selected with 1 mg/mL of G418 (Coolaber, China) at 48 h after transfection. After two weeks of selection, the expression level of ACE2 was determined by immunoblotting.

### SARS-CoV-2 Spike protein pseudovirus generation and titration assay

A recombinant replication-deficient vesicular stomatitis virus (VSV) vector that encodes SARS-CoV-2 Spike protein instead of the VSV-glycoprotein (VSV-G) is pseudotyped with SARS-CoV-2 spike (S) derived from SARS-CoV-2 Wuhan-Hu-1 strain (GeneBank: YP_009724390.1) (Li *et al*, 2020), which is a kind gift from Prof Aihua Zheng, Institute of Zoology, Chinese Academy of Sciences. The pseudovirus was harvested and passaged on Vero E6 cells. Briefly, The Vero E6 cells were cultured in T75 flasks until reaching 90% confluence. The medium was replaced by serum-free DMEM supplemented with 5 μg/mL polybrene (solarbio, China) and pseudovirus (MOI=0.1) was added into cells. After 8 h infection, the medium was replaced by 20 mL of DMEM with 2% (*v/v*) FBS and 1% (*w/v*) HEPES. After 72 h incubation, three freeze/thaw cycles were performed to release pseudovirus from cells. The pseudovirus-containing culture supernatant was harvested and centrifuged at 800 g for 10 min and filtered by ultrafiltration centrifuge (50 K, Millipore). The concentrated pseudovirus was filtered (0.22 µm pore size; Biosharp, China) and stored at −80 ℃ until use.

The expression of S protein in infected cells and culture supernatant was confirmed by Western blotting. The titer of SARS-CoV-2 pseudovirus was determined by microdose CPE assay. The Vero E6 cells (1×10^4^ cells/well) were seeded into 96-well plates one day prior to infection. The pseudovirus was serially diluted ten-fold in serum-free DMEM and added to plates. Following 3 h incubation, the culture media was replaced by DMEM containing 2% (*v/v*) FBS and cultured at 37℃ with 5% CO2 for 48 h. The cells were checked under a microscope for the presence of CPE every day. The 50% tissue culture infectious dose (TCID50) was calculated according to the Reed-Muench method.

### Immunoblotting

The cell lysis and concentrated supernatant were prepared in loading buffer and separated by 6% (for S protein and ACE2) or 10% (for GAPDH) SDS-PAGE. The primary antibodies used were a mouse monoclonal antibody to ACE2 (1:2000 dilution, Proteintech, China), a rabbit polyclonal antibody to SARS-CoV-2 Spike protein (1:2000 dilution, Abclonal, China), a mouse monoclonal antibody to GAPDH (1:3000 dilution, Abways, China), and their corresponding secondary antibodies (1:5000 dilution, Abways). The signal was detected with Enhanced ECL Chemiluminescent Substrate Kit (YEASEN, China) and visualized with the ChemiSciope 6100 Imaging System (CLINX, China).

### Pseudovirus-based CPE inhibition assay

The Vero E6 cells were seeded into 96-well plate at 1×10^4^ cells per well 24 h before infection. The isolated glycoproteins, BSA, and raw milk proteins were serially diluted with serum-free DMEM and pre-incubated with pseudovirus (31.6 TCID50 per well) at 37°C for 1 h, respectively. The proteins-pseudovirus mixture was added to plates. Three hours later, the cells were washed with sterile PBS and cultured with DMEM containing 2% (*v/v*) FBS for 48 h. The CPE was determined by a microscope, and the half maximal inhibitory concentration (IC50) of isolated glycoproteins was calculated with non-linear regression, i.e. log (concentration of inhibitor) vs response (four parameters), using GraphPad Prism 7 (GraphPad Software, Inc., San Diego, CA, USA) from three independent repeated experiments.

### Plaque inhibition assay

Vero E6 cells were seeded in six-well plates at 4 × 10^5^ cells/well in triplicate, one day before infection. Pseudovirus (94.8 TCID50 per well) pre-incubated with isolated glycoproteins, BSA, and raw milk proteins (final concentration is 0.13 μg/μL) at 37 °C for 1 h, respectively. The mixtures were incubated with cells in 96-well plates for 3 h. The cells incubated with or without pseudovirus served as positive and negative controls. After that, the cells were washed with PBS and overlaid with DMEM-Avicel solution (DMEM with 2% (*v/v*) FBS and 1% (*w/v*) Avicel) for 72 h at 37°C with 5% CO2. The cells were fixed in 4% paraformaldehyde (YEASEN) and strained in 1% crystal violet staining solution (Coolaber, China). The number of plaques of each group was counted from triplicate wells.

### Immunofluorescence assay

The effect of isolated glycoproteins on the binding of pseudovirus to hACE2/293A cells was analyzed by immunofluorescence assay. Briefly, the hACE2/293A cells (3 × 10^4^ cells) were inoculated into 30 mm confocal culture dishes (NEST, China) 24 h before infection. Pseudovirus (63 TCID50 per dish) pre-incubated with isolated glycoproteins, BSA, and raw milk proteins (final concentration is 0.13 μg/μL) at 37 °C for 1 h, and then, the mixtures were further incubated with cells for 6 h at 37 °C. After washing with PBS, the cells were immobilized by 0.2% Triton X-100 in 4% paraformaldehyde for 30 min and blocked by 5% BSA in PBS solution for 1 h at room temperature. Following, the cells were incubated with the primary antibody of S protein diluted at 1:200 in PBS with 3% BSA overnight at 4°C, and then incubated with Cy3 labeled second antibody (dilution: 1:500, Bioss, China) for 1 h at room temperature in darkness. The nucleus was stained with 4,6-Diamidino-2-phenylindole (DAPI, 1 µg/mL in PBS) (Roche; Basel, Switzerland). Finally, the Laser Scanning Confocal Microscope (TCS SP8, Leica, Germany) was used to acquire the images with the merge channels of Cy3 and DAPI at the same exposure time and shown on the same scale.

### Statistical analysis

Descriptive statistics and non-linear regression using GraphPad Prism (version 7). Comparisons between groups were analyzed using Mann-Whitney tests or one-way analysis of variance (ANOVA; Tukey’s multiple comparisons test), respectively. P values less than 0.05 was considered to be statistically significant.

## Acknowledgments

We are grateful to the National Natural Science Foundation of China (Grant No. 32101030), the China Postdoctoral Science Foundation (Grant No. 2020M673628XB), the Natural Science Foundation of Shaanxi Province (Grant No.2021JM-319 and 2021JQ-446), and Shaanxi Province Key R&D Program (No. 2021ZDLSF01-04) for financial support.

## Author contributions

H.Y., W.C., and Z.L. conceived the project. H.Y., W.C., J.S., X.W., X.B., J.Q., D.W., H.C., and X.L.W. carried out experiments. All authors analyzed and discussed the data. H.Y., W.C. and Z.L. wrote the manuscript. All authors read and approved the manuscript.

## Data availability

This study includes no data deposited in external repositories.

## Expanded View Figure legends

**Figure EV1. Molecular dynamics simulated the interaction of glycans and GRDs.** The “complex” type N-glycans were added at N-glycosites of N53, N90, N103, N322, and N546 of ACE2, N343 of RBD, N165, and N234 of NTD on S1 subunit. The distances of the terminal of glycans on these sites and three GRDs of ACE and S1 subunit were monitored during a 100 ns MD simulation. As a result, the average distances of N546-GRD1, N322-GRD2 and N53-GRD2 fluctuate were approximately 15 Å, and the distances of N343-GRD3 and N165-GRD3 fluctuated between 20 to 35 Å.

**Figure EV2. Detection and evaluation of SARS-CoV-2 pseudovirus in infected cells.**

A, B The expression level of S protein in pseudovirus-infected cells and culture supernatant. After 72 hours post-infection, the cell lysate of Vero E6 cells (A) and culture supernatant (B) were analyzed by western blot using an antibody against S protein.

C CPE and plaques were observed in Vero E6 cells infected with SARS-CoV-2 pseudovirus. Vero E6 cells were cultured in six-well plates (4 × 105 cells per well) and then infected with pseudovirus (94.8 TCID50 per well) for 3 h. After washed with PBS, the cells were cultured in DMEM containing 2% FBS and 2% FBS/1% avicel DMEM medium, respectively. The dramatic CPE (upper right) and visible plaques (low right) in Vero E6 cells were observed after 72 h post-infection.

D Overexpression ACE2 in 293A cells. The 293A cells were transfected with pCDNA3.1-ACE2 (human)-3×FLAG plasmid and selected with G418 (1 mg/mL) for 2 weeks. The expression level of ACE2 in 293A and 293A-hACE2 cells were analyzed by western blot.

